# Searching for biological feedstock material: 3D printing of wood particles from house borer and drywood termite frass

**DOI:** 10.1101/2020.05.27.118562

**Authors:** Rudy Plarre, Andrea Zocca, Andrea Spitzer, Sigrid Benemann, Anna A. Gorbushina, Yuexuan Li, Janka Wilbig, Jens Günste

## Abstract

Frass (fine powdery refuse or fragile perforated wood produced by the activity of boring insects) of larvae of the European house borer and of drywood termites was tested as a natural and novel feedstock for 3D-printing of wood-based materials. Small particles produced by the drywood termite *Incisitermes marginipennis* and the European house borer (EHB) *Hylotrupes bajulus* during feeding in construction timber, were used. Frass is a powdery material of particularly consistent quality that is essentially biologically processed wood mixed with debris of wood and faeces. The filigree-like particles flow easily permitting the build-up of wood-based structures in a layer wise fashion using the Binder Jetting printing process. The quality of powders produced by different insect species was compared along with the processing steps and properties of the printed parts. Drywood termite frass with a HR = 1.1 with ρBulk = 0.67 g.cm^-3^ and ρTap = 0.74 g.cm^-3^ was perfectly suited to deposition of uniformly packed layers in 3D printing. We suggest that a variety of naturally available feedstocks could be used in environmentally responsible approaches to scientific material sciences/additive manufacturing.

## 1. Introduction

Anthropogenic perturbations of natural ecosystems are omnipresent: materials and products of human activity are superimposed on natural cycles everywhere. According to Schellnhuber 1999 [1] there are two main components: the ecosphere N and the human factor H. N consists of intricate linkages between the atmosphere, hydrosphere, cryosphere, lithosphere, biosphere, etc, while the human factor H aggregates all actions and products along with a metaphysical component of human activity.

Sustainable coevolution of the ecosphere and the anthroposphere requires fresh scientific attitudes and approaches, including completely new ways of manufacturing. The ever-increasing human impact on the planet requires the deliberate coupling of natural feedstocks to novel manufacturing process. This way the life cycle of the products and materials can be determined early in production. Substituting dedicated feedstocks for additive manufacturing (AM) with surplus natural materials is one way to substantially increase AM sustainability while concomitantly providing high-value outputs for “pre-owned” materials/products.

As a general rule for all production processes, natural and recycled feedstocks should take preference to dedicated ones – especially in the context of a circular economy. Deliberately developing naturally available feedstock constitutes an environmentally responsible scientific approach in material sciences.

### 1.1 Additive manufacturing

Adding material to form an object instead of subtracting material from an excessively large block is a new trend in manufacturing technologies which is currently stimulating an entire industry [2]. In the majority of additive manufacturing processes, the material is added layer by layer. The raw material (feedstock) is fed into the process as a powder/granulate, paste or suspension, as it is in a state optimized for the layer deposition process. In the manufacturing process itself, the feedstock is used to build up the desired object and it is simultaneously transferred into a state possessing its final physical properties, or at least providing enough mechanical strength to transfer the configured object to further processing steps. Adding instead of subtracting material implies more than just flexibility in design. Multi material processing, the generation of unique properties and functionalities and functionally graded materials are just some facets intrinsic to AM. One of the most popular and widespread additive manufacturing technologies, the “Binder Jetting” (BJ), is making use of a powdery material as feedstock [3, 4], see also Fig 1. A layer of powdered material is first spread as a layer and subsequently the corresponding layer information of the object manufactured is selectively inscribed by a printing head, spraying individual droplets of a binding liquid onto the powder layer, thus selectively consolidating the powder and defining the cross-section of the object in a respective layer.

**Fig 1.**
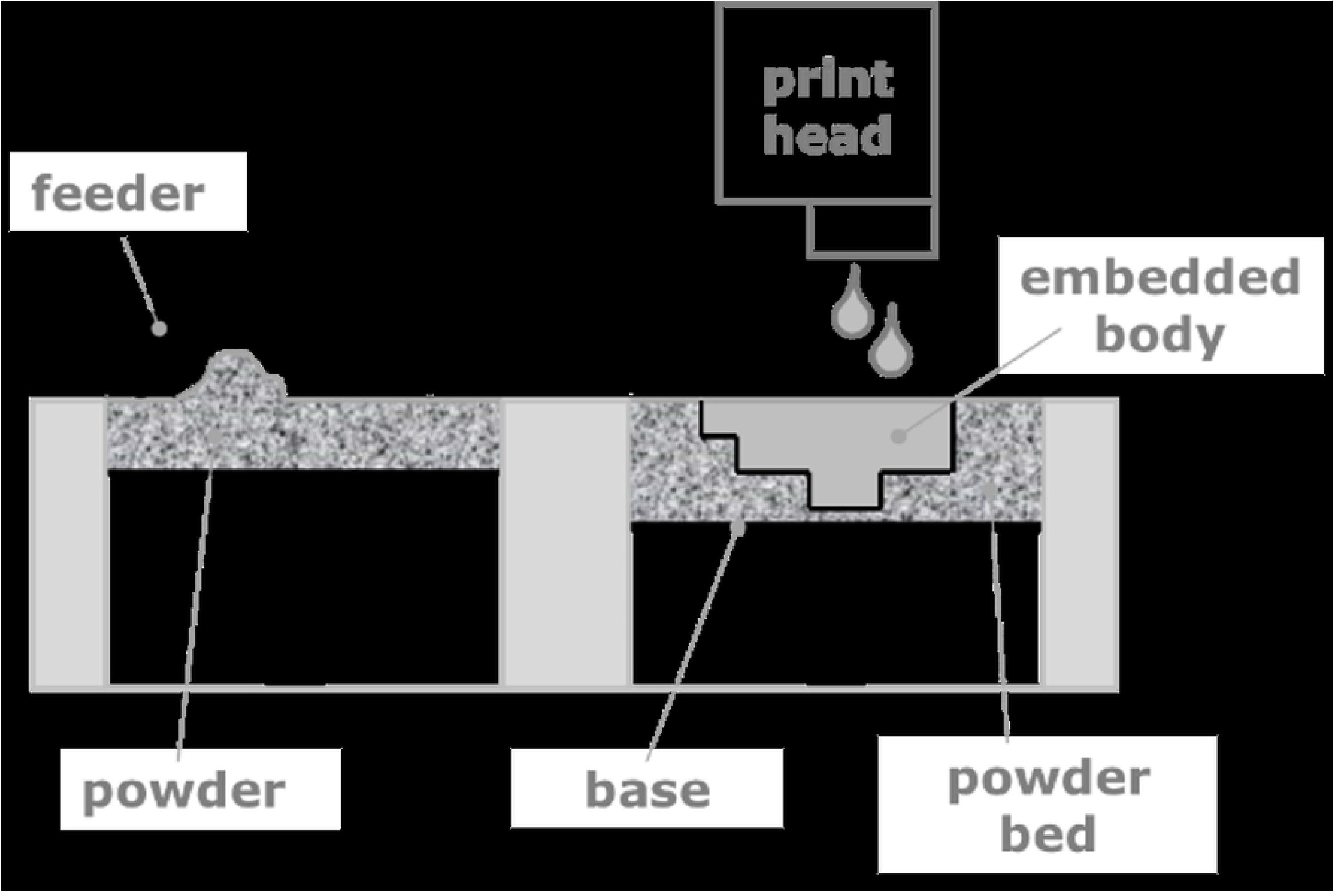
Schematic of the Binder Jetting (BJ) process.

The popularity of BJ is based on the fact, that it virtually can accept all powdery materials which provide sufficient flowability to be spread as a homogeneous thin layer. This flexibility has stimulated the creativity of many research groups envisioning the use of abundant materials, such as sand [5,6,7], or recyclable material in powdery form for upgrading waste for the manufacture of new products [8]. Costs and resources for the initial material synthesis can be saved. However, the refinement of powdery raw materials to powders well suited for BJ remains a mandatory processing step if the recycled material is not directly obtained in the appropriate powdery state [9].

Here a strategy of coupling the development of the most promising AM approaches to considerations of the use of natural or Nature-recycled materials as feedstocks was applied. In the present work the powdery material which remains when house borer larvae or drywood termites feed on wood, the so called frass, is used as a novel feedstock. 3D printing of wood chops [10–14] or plastic-wood composites has already been proven to be a feasible way for obtaining objects with wooden haptics. Dedicated feedstocks have been processed from wood and have been adapted to the respective printing process by refining it with polymeric additives [15–18]. In terms of sustainability, 3D printing of wood-based materials from house borer and termite frass is going one step further as it is not only using naturally occurring but also naturally processed materials directly as feedstocks.

In the BJ process, the binder system, which is used to consolidate the powdery material to form an object, has to fulfill multiple requirements. As a liquid, it has to be of an appropriate viscosity and surface tension to be dosed by a commercial printer head [4, 19, 20]. In order to penetrate the deposited layer, it must also moisten the powdery feedstock. Moreover, it ideally interacts with the powdery material to form a strong interparticle adhesion. In most cases, addition of a binder does not result in a significant densification of the powder. Hence, BJ is typically providing porous parts, which may be densified by sintering/melting, infiltration of additional material or post compaction by isostatic pressing etc. [21, 22].

### 1.2 Feedstocks from timber

A major interest exists in 3D printing for ecofriendly and recycled materials, particularly in Binder Jetting [11–16, 23, 24]. BJ can utilize basically anything that can be powdered to an appropriate particle size. The particle size of the powders is essential for obtaining sufficient flowability for the deposition of defect-free layers: In case of too fine powders, the flowability will be poor, in case of too coarse-grained powders the definition of the part will be imprecise [25]. Wood particles may be obtained as byproducts from wood machining, such as saw dust, or are deliberately processed from wood. For obtaining suitable feedstocks for 3D printing, in most published works the wood particles are mixed into a polymer or mixed with other binning phases [15, 17, 26–29]. On the other hand, timber can also be naturally processed into printable powders by e. g. insects feeding on wood. We have used small particles from feeding byproducts of the European house borer *Hylotrupes bajulus* and drywood termite *Incisitermes marginipennis* as raw material for 3D-printing.

Insects in construction timber share several anatomic and physiological features which make them to appear perfectly adapted to this environment. Wood is an inhomogeneous poriferous matrix containing mainly cellulose and lignin. Their relative amounts vary between heartwood and sapwood as well as in the early and late wood of the annual rings. This results in local strength differences and an uneven distribution of essential nutrients for the insects. The larvae of wood boring beetles or drywood termites have strong mandibles which allow abrasion of all parts. The cellulose is the main hydrocarbon source usually digested with the aid of cellulase-producing microorganisms. However, the rare nitrogen containing elements of wood are the limiting factors. In order to access as much nitrogen as possible much more wood is consumed by wood feeding insects than actually needed for development. Larvae of *H. bajulus*, e. g. excavate throughout the sapwood leaving extensive tunnels filled with frass. The frass contains either loosely chopped off wood particle (debris) or dense-packed faeces. The latter is of cylindrical shape made up out of himidigested cellulose/lignin conglomerates. Destructions of insects feeding on build-in timber can cause severe danger and precautions are needed. Laboratories like BAM therefore rear large pest populations to test different control strategies for efficacy evaluation in pest control. The rearing byproducts like the frass were of no further use and discarded. However, after being modified by the insect digestive system the former non-uniform wooden material is changed to a homogeneous compact cellulose-lignin mixture and becomes suitable for further technical applications, such as 3D printing, without any further processing. In the present study we have evaluated, the so called frass, as feedstocks for 3D printing. The morphology of the drywood termite frass is quite different to house borer frass. While drywood termite frass appears as six-sided pellets almost uniform in size the frass from house borer is sawdust-like and more irregularly shaped. In contrary to drywood termites, the house borer larvae digest only part of the abraded wood with their frass containing loosely chopped off debris as well as more dense-packed faeces. As will be pointed out further down, the 3D printing of frass as a feedstock is feasible and printed parts can be further processed, e.g., by pyrolysis into carbon and subsequent to reactive silicon infiltration, which may serve as a template for SiC ceramics featuring unique structural properties.

## 2. Materials and methods

Larvae of EHB were reared at constant conditions of 28 ±2 °C and 75 ± 5 % r.h. During the first days after hatching from the eggs, larvae were manually inserted into pine, *Pinus sylvestris*, sapwood blocks (1.5 × 2.5 × 5 cm^3^) enriched with peptone and yeast. This enrichment with nutrients was carried out by impregnating the sapwood with an aqueous solution of 1 % peptone and 0.3 % yeast at low pressure of 100 to 200 mbar for 30 minutes to speed up development. Two larvae per wood block were allowed to feed for approximately six months before being individually transferred into pure pine sapwood blocks (3 × 4 × 5.5 cm3), not enriched with any nutrients. While feeding in the wood, the larvae produce debris and faeces which are left behind in the frass tunnels. Debris is undigested wooden material derived from abrasion processes when the larvae’s mandibles carve on the wood. It usually bypasses the larvae during movement through the wood. While debris are of undefined structure, the faeces are densely packed into cylindrical pellets when leaving the larva’s hindgut. As the wood is increasingly consumed by the larvae, frass (debris and faeces) eventually trickle out and can be collected in larger amounts.

Using a vibratory sieve shaker (Analysettre 3 spartan, Fritsch, 55743 Idar-Oberstein, Germany) EHB frass was sieved for 30 min at 1 mm amplitude. A fraction with particle size distribution of 45 μm – 100 μm, which amounts to 17% of the total frass, 57% above 100 μm and 26% below 45 μm, was then used for 3D-printing, see also Fig 2. The frass particles show a considerable flowability although the particle shape is rather flake-like than spherical. The Hausner ratio (HR), as a measure of flowability [29], was determined according to the relation shown in **Error! Reference source not found.**:

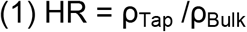

where HR is the Hausner ratio, ρ_Bulk_ displays the freely settled bulk density of the powder and ρ_Tap_ the bulk density after a given number of tapping cycles, at which the bulk density is in a plateau, in g/cm^3^ With ρ_Bulk_ = 0,14 g/cm^3^ and ρ_Tap_ = 0,18 g/cm^3^, the EHB frass particles have a Hausner ratio of 1.25 which displays a fair flowability according to the classification [30].

**Fig 2.**
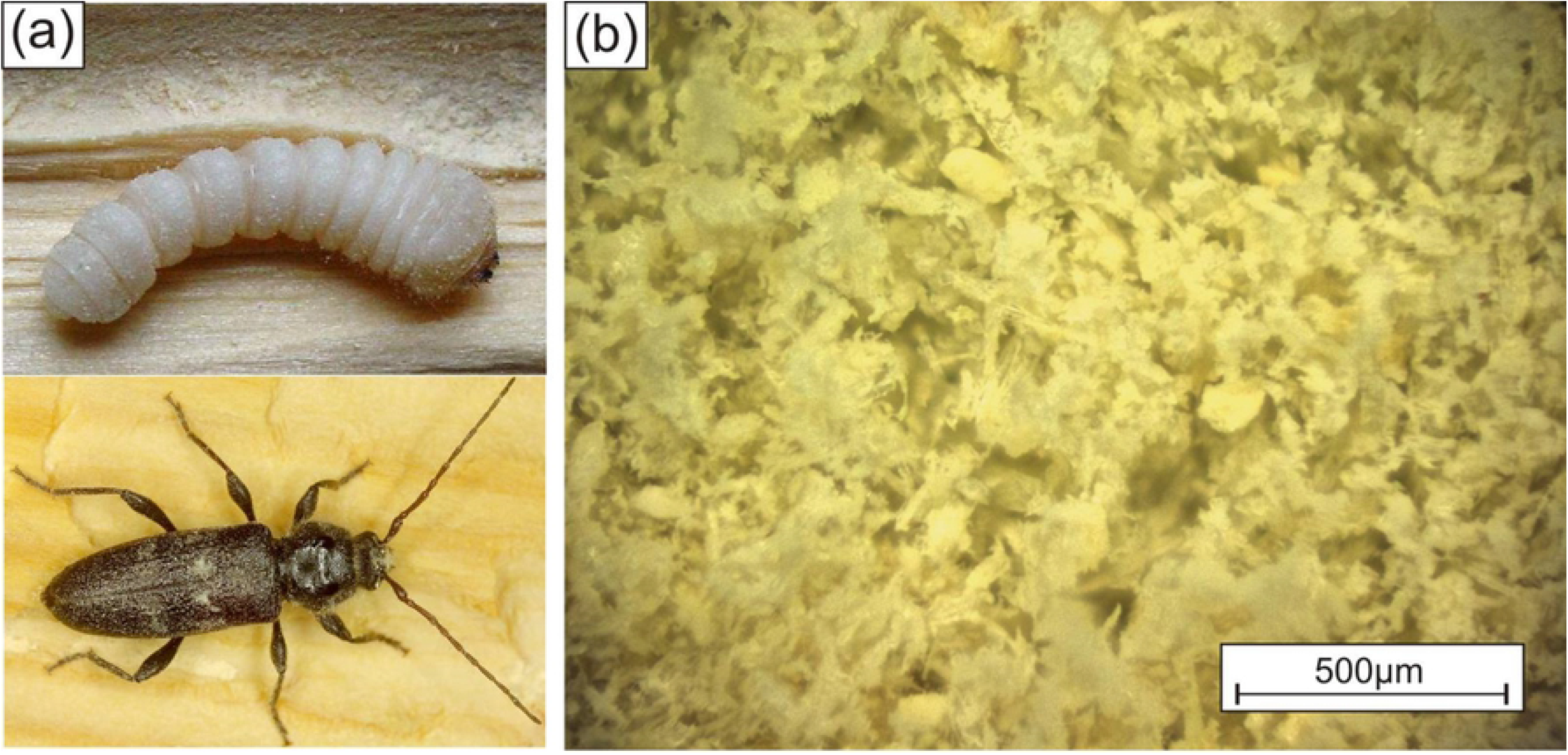
a) European house borer (Hylotrupes bajulus) full grown larva (top) and adult beetle (bottom); b) the sieved frass (in a particle size fraction of 45 to 100 μm) produced by larvae and used for 3D-printing.

Within this study, frass from drywood termites has been considered as feedstock for 3D printing, as well. Termites rely on fungi, protists and bacteria that live in their gut to break down the wood and digest lignin and cellulose. In comparison to the feeding byproducts of EHB the drywood frass contains six-sided pellets almost uniform in size and reveal an excellent flowability required for the layer wise buildup of wooden structures, see also Fig 3. The pellets are very compact and composed of fine fibers and particles.

**Fig 3.**
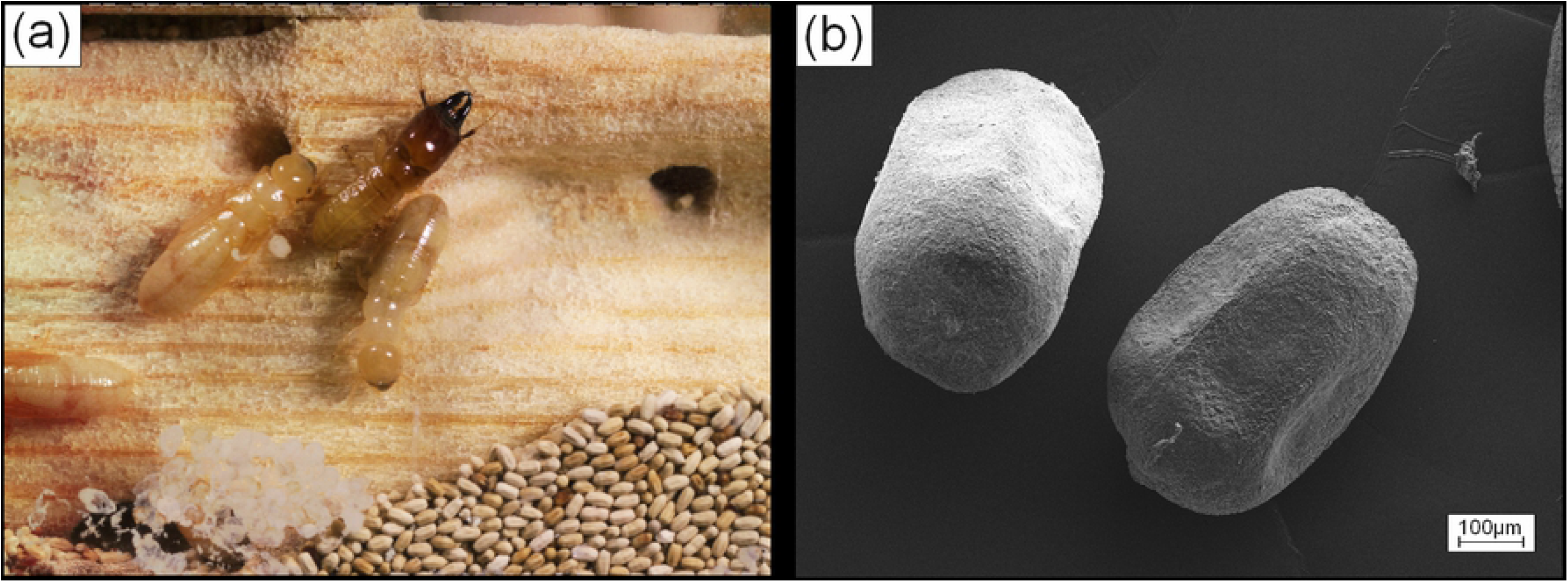
a) Drywood termites (Incisitermes marginipennis), soldier and worker with frass; b) SEM micrograph of single pellets of drywood termite frass.

Their good flowability, HR = 1.1 with ρ_Bulk_ = 0,67 g/cm^3^ and ρ_Tap_ = 0.74 g/cm^3^, making them perfectly suited for the deposition of uniformly packed layers in 3D printing. One general problem with compacted porous particles, i.e., the frass pellets, however, exists in the binder jetting process: Driven by capillary forces, the binder is drawn into the fine network of pores within the individual pellets. Primarily, all capillaries within the pellets have to be filled before a sufficient amount of binder remains available for gluing the pellets to each other effectively. In the competition among capillaries with smaller diameters within individual pellets and capillaries with larger diameters between individual pellets, the smaller diameters are providing a stronger drag force for the binder uptake. Hence, a lot of binder is consumed before excessive binder is available for interparticle gluing. Unfortunately, due to the binder uptake the pellets are not swelling and do not lose their structural integrity, which could help to disintegrate and glue them together. Therefore, in the printing process a high binder saturation of the powder bed is required for consolidating a structure while an excessive amount of binder stays in the pellets.

## 3. Results and discussion

### 3.1 Powder-based 3D-printing by Binder Jetting (BJ) of EHB frass

Due to the low packing density of the ESH frass, parts obtained from 3D printing have shown low mechanical strength and were considered as not useful in the as-printed state. On the other hand, they show a considerably well detail quality and could serve as preforms which are converted into ceramic structures as initially introduced by a concept from Greil et al. [31].

Using a commercial 3D printing machine (RX-1, Prometal RCT GmbH, Augsburg, Germany) cubic specimens with dimensions of 9 mm3 and rectangular struts were printed. The binder used was a water based commercial system provided by ExOne GmbH, Augsburg Germany (PM-B-SR2-02, viscosity of 10.7 mPa·s @ 1000s-1). Cross-linking of the binder was carried out after each layer printing in the thermal curing station of the printer. The precision of the printed geometry depended on several processing parameters (e. g. powder particle size, flowability, and layer thickness) as well as binder saturation. Printing parameters such as binder saturation, layer thickness and curing time were varied. The binder saturation is a parameter useful to evaluate the amount of binder used to glue a certain quantity of powder, because it gives the ratio between the volume of binder spread out in a volumetric unit (voxel) and the free volume, not filled with powder, in the same voxel. A saturation S of 100%, that means all porosity of the powder bed is initially filled by binder, was chosen for printing EHB frass. The layer thickness which turned out to be most appropriate for a reproducible deposition of uniform layers was 100μm.

A structure printed according to the model from Fig 4 is shown in Fig 5a. The model structure is reproduced very well except for the right-handed side of the cube, which was the bottom plane during the printing process. Clearly a convex instead of planar shape can be recognized. This distortion arises from an oversaturation with binder. Also indicative for an oversaturation is the reduction in the diameter of the printed capillaries. Instead of the designed square shaped cross section of 1mm edge length, the edge length is reduced to approximately 500 μm.

**Fig 4.**
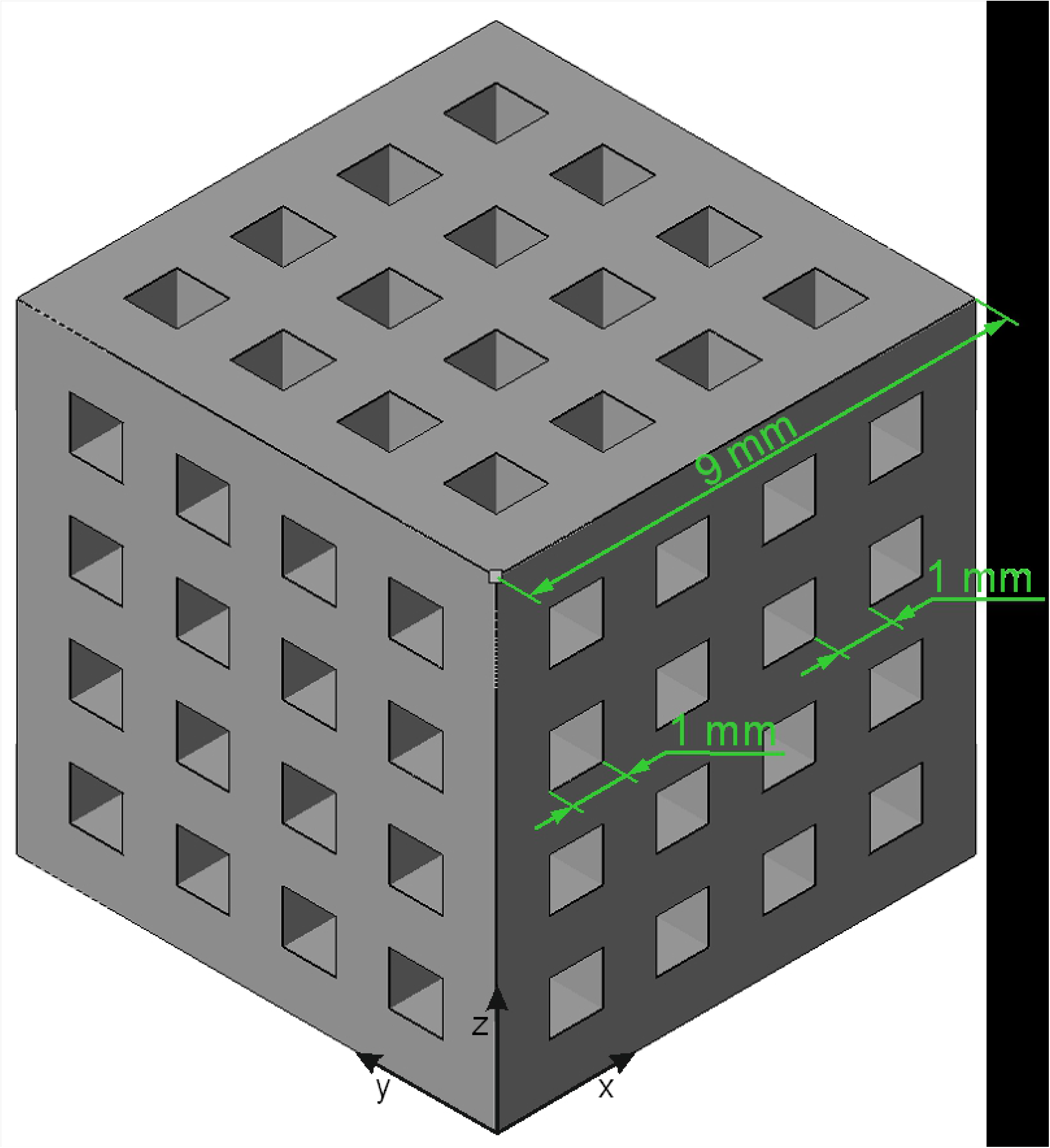
Schematic drawing, including dimensions, of the specimen printed from EHB frass.

**Fig 5.**
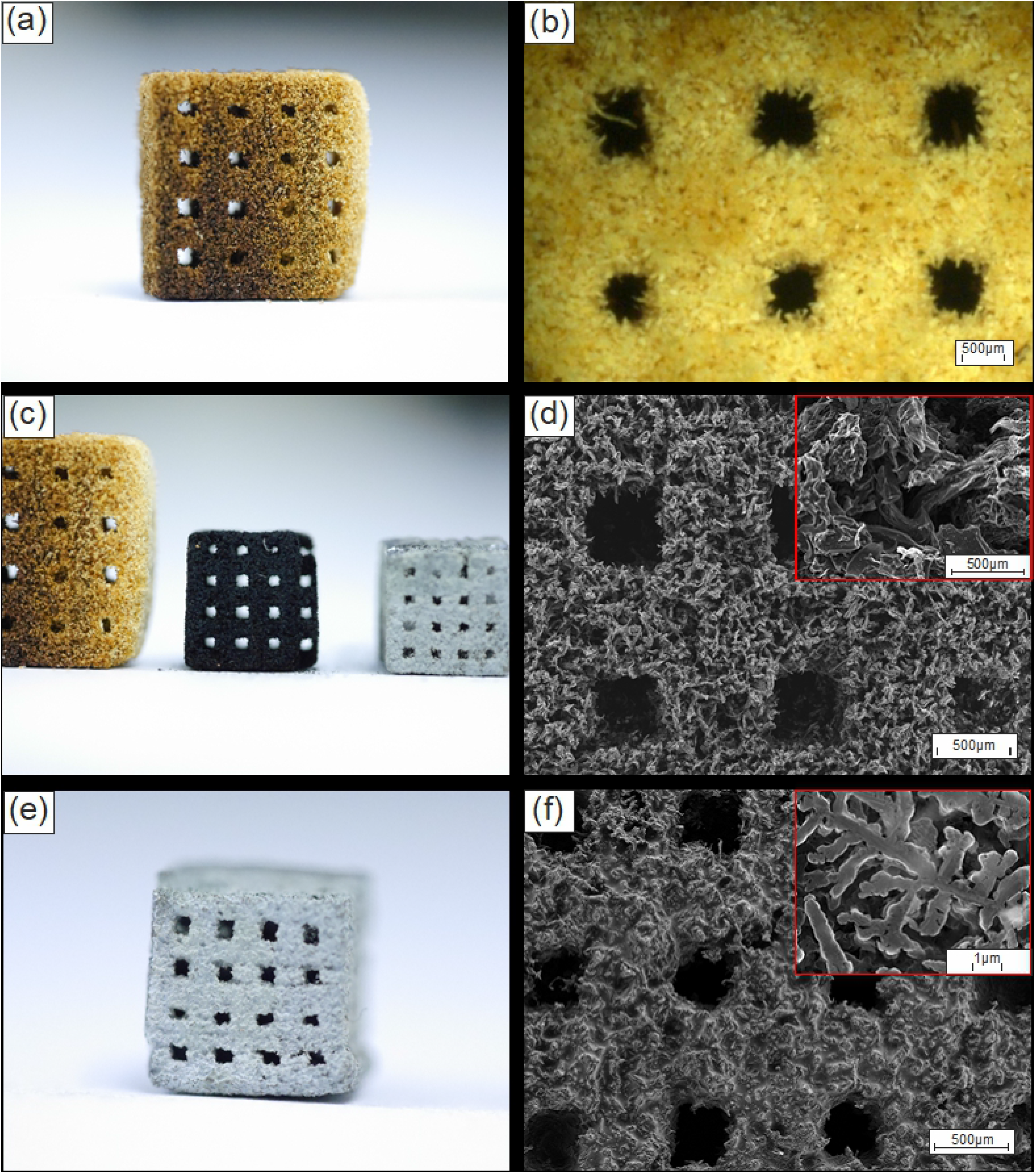
Imaging of the manufactured specimens from European house borer (EHB) frass. a) Photograph of a printed cubic sample. b) Light microscope image of the sample from a), showing the wooden chops and macroscopic channels. c) Photograph of a pyrolized specimen (middle, black). d) SEM-image of the pyrolized part. Individual chops of the frass are still observable. e) Photograph of the Si-infiltrated sample. f) SEM-image of the infiltrated sample, the remained unreacted silicon forms dendritic structures as can be seen in the inset.

The density of the printed parts is very low, but sufficient for transferring them to additional process steps such as pyrolysis and silicon infiltration. The frass particle were hardly compacted during layer deposition. Addition of binder did not result in a densification. Fig 5b shows details of the fluffy structure of the as-printed part. In order to enhance the mechanical strength, the printed part was pyrolyzed in a vacuum furnace at 1400 °C resulting in a structure entirely made from carbon (see also Fig 5c). Due to a significant loss of volatile fractions during pyrolysis, the structure shrank by approximately 40% (linear shrinkage). Nonetheless, due to a homogeneous shrinkage, no additional defects are generated. According to a strategy applied to natural cellulose containing materials such as wood [31], this carbon preform can be infiltrated by silicon to form silicon carbide. In an additional heat treatment at about 1470 °C, the carbon preform it brought into contact with liquid silicon. Due to an excessive supply of silicon, the structure is also filled with metallic silicon and a silicon infiltrated silicon carbide structure is formed. The microstructures of the pyrolyzed graphitic and the infiltrated parts are shown in Figs 5d and 5f, respectively.

Fig 5d reveals the flaky structure of the pyrolyzed wood chops. During silicon infiltration, these carbon flakes are acting as wicks for the liquid silicon, which viscosity is at 1470 °C comparable or even lower than water [32], resulting in a dendritic structure of the infiltrated material, Fig 5f.

As the mechanical and structural properties of the silicon infiltrated parts critically depend on the microstructure of the 3D printed preform, a discussion of possible applications is beyond the scope of the present study. Nevertheless, it could be shown, that 3D printing of EHB frass is not only possible but yields parts which can be further processed.

### 3.2 Powder-based 3D-printing by Binder Jetting (BJ) of drywood termite frass

Frass from dry wood termites reveals an excellent flowability and, thus, relatively high packing densities could be obtained during layer deposition. The approximately four times higher packing density, as compared to the EHB frass, appears promising for even load bearing applications of printed parts, without further processing. On the other hand, the size of the individual pellets in the range of a few 100 μm is unusual large for BJ. In order to adapt the printing process to the large size of the pellets (ca. 500 μm on their long axis), layer thicknesses of minimum 600 μm have been evaluated and 800 μm was found optimal for the deposition of smooth layers. In order to compensate the binder uptake of the single pellets, see materials and methods section, a binder saturation of 166% was required. It was found difficult to deposit layers on top of the first printed cross sections, due to adhesion of the binder saturated pellets to the recoater. This problem could be solved by applying a gas flow through the powder bed, providing an additional force stabilizing already deposited material. This technology has been introduced recently for the deposition of powders with poor flowability [33]. Figure 7 shows a part printed from termite frass according to the model shown in Fig 6.

**Fig 6.**
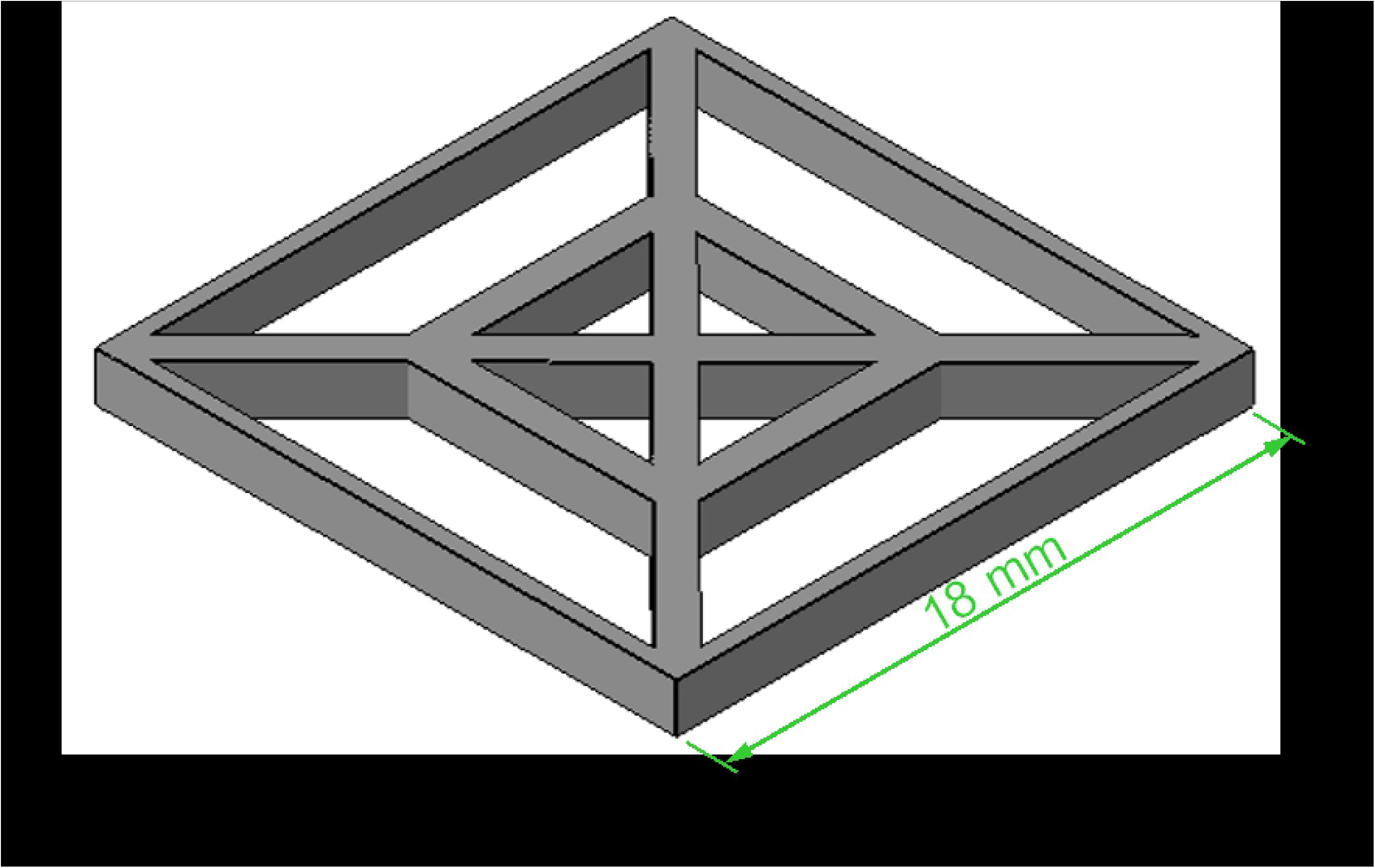
Schematic drawing, including dimensions, of the specimen printed from drywood termite frass.

**Fig 7.**
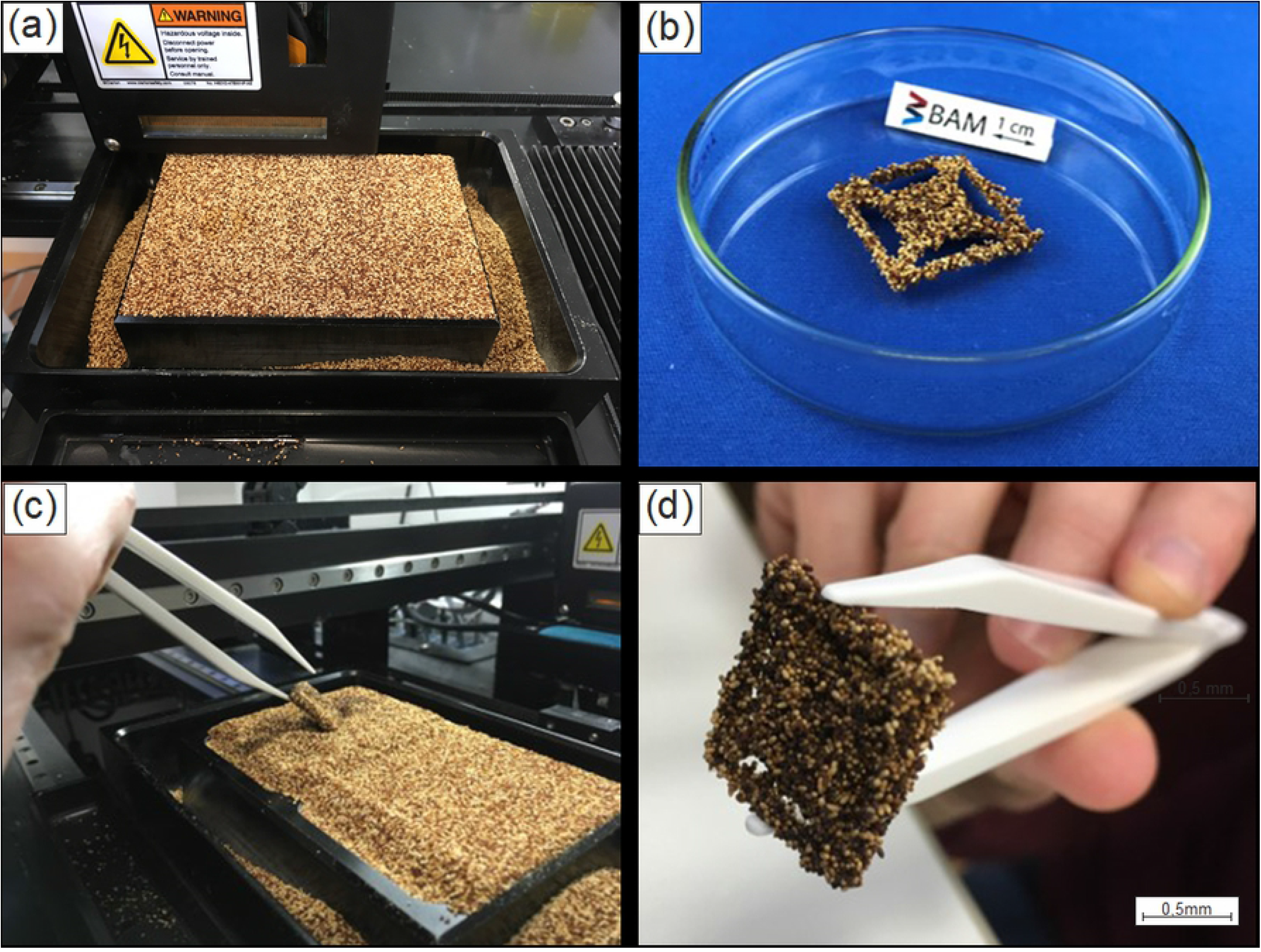
a), Imaging of the BJ powder bed filled with drywood termite frass, upper left corner of the image the print head is visible. b) part printed according to figure 6. c) part from d) as it is taken out of the powder bed after printing. d) part printed with same geometry as shown in b) but reduced in size by a scaling factor of 0.5.

In order to get an impression about the possibility to reproduce fine structural features in BJ with particles as large as 500 μm, the model structure from Fig 6 has been printed in two sizes, original size, as indicated in the figure, and reduced in size by a scaling factor of 0.5.

With a size of approximately 300 μm at the short and 500 μm at the long axis individual pellets of the termite frass are almost one order of magnitude larger than powders typically used in 3D printing. However, for printing wooden objects like furniture [16], coarse grained powders are beneficial for the deposition of thicker layers and higher building rates could be achieved. Structural details are reproduced on a scale of several millimeters but definitely not below one millimeter, compare Figs 7b and d.

## 4. Conclusion

Here we considered wood processed by the drywood termites *Incisitermes marginipennis* and the European house borer (EHB) *Hylotrupes bajulus* as feedstocks for 3D printing. This approach follows the general strategy of developing naturally available feedstocks as environmentally responsible substrates in material sciences. The quality of the powdery feedstocks, the so-called frass provided by these termites during feeding in construction timber was very different in terms of process ability in 3D printing. EHB frass lead to a flaky structure with poor packing density, whereas the termite frass consisting of pellets of almost uniform size and packed very well. Despite of the different packing densities, both feedstocks could be spread out into thin homogeneous layers for the build-up of structures in the Binder jetting 3D printing process. At approximately 300 μm along the short- and 500 μm at the long-axis, pellets of termite frass are almost an order of magnitude larger than powders typically used in 3D printing. In printing wooden objects like furniture, coarse-grained powders allow the deposition of thicker layers and higher construction rates. The fine, sawdust-like frass particles of EHB flow especially well allowing deposition of reproducible layers and, because of their smaller particle size, are well suited to the printing of filigree structures.

